# Jaeger: an accurate and fast deep-learning tool to detect bacteriophage sequences

**DOI:** 10.1101/2024.09.24.612722

**Authors:** Yasas Wijesekara, Ling-Yi Wu, Rick Beeloo, Piotr Rozwalak, Ernestina Hauptfeld, Swapnil P. Doijad, Bas E. Dutilh, Lars Kaderali

## Abstract

Viruses are integral to every biome on Earth, yet we still need a more comprehensive picture of their identity and global distribution. Global metagenomics sequencing efforts revealed the genomic content of tens of thousands of environmental samples, however identifying the viral sequences in these datasets remains challenging due to their vast genomic diversity. Here, we address identifying bacteriophage sequences in unlabeled sequencing data. In a recent benchmarking paper, we observed that existing deep-learning tools show a high true positive rate, but may also produce many false positives when confronted with divergent sequences. To tackle this challenge, we introduce Jaeger, a novel deep-learning method designed specifically for identifying bacteriophage genome fragments. Extensive benchmarking on the IMG/VR database and real-world metagenomes reveals Jaeger’s consistent high sensitivity (0.87) and precision (0.92). Applying Jaeger to over 16,000 metagenomic assemblies from the MGnify database yielded over five million putative phage contigs. On average, Jaeger is around 20 times faster than the other state-of-the-art methods. Jaeger is available at https://github.com/MGXlab/Jaeger.

## Introduction

Viruses are the most abundant life forms on the Earth and infect members of all three domains of cellular life ^1^. Most of these viruses target bacterial or archaeal hosts, we refer to these as phages. Phages can affect microbial populations by reprogramming metabolic pathways ^2^, inducing phenotypic changes ^3^, and influencing nutrient cycling ^4,5^. Despite their wide distribution, from the mammalian gut ^6–8^ and the rhizosphere ^9,10^ to the ocean depths ^11,12^, the identification of phages remained limited before metagenomics ^6–8^. Metagenomic sequence assemblies often yield contigs originating from diverse organisms, including eukaryotes, bacteria, archaea, and viruses. Identifying viral sequences, especially phages, in such data poses challenges for several reasons. First, short contigs may lack sufficient phage-specific signals for identification ^13^. Second, even with long contigs, homology-based identification may be challenging as many newly detected phages lack similarity to known ones in the databases ^14^. Third, there are no universal marker genes to identify phages ^15,16^. Lastly, distinguishing bacterial and phage contigs is complicated by phage-like regions on bacterial contigs ^17–19^. Hence, many novel phages remain concealed within metagenomic datasets.

Phages have traditionally been discovered by comparing sequences from metagenome assemblies with those in databases. Bioinformatics pipelines incorporating tools such as the BLAST suite ^20^ and Diamond ^21,22^ have been used for this purpose ^23–26^. However, these methods may not identify phages that differ significantly from those in the databases. To address this limitation, Hidden Markov Model (HMM) databases of phage protein families, paired with tools like HHSuite ^27^ or HMMER ^28^, have been employed due to their high sensitivity.

Despite their effectiveness, these methods face challenges. Selecting suitable sequence similarity cut-offs is crucial; a higher cut-off improves precision but reduces True Positive Rate (TPR), and vice versa ^29^. Furthermore, the large size of sequence databases poses difficulties in terms of computational resources and processing time required by these approaches.

Later, several supervised learning methods were introduced to enhance virus identification by integrating sequence database search results with additional genomic attributes, such as average gene length, gene orientation, nucleotide k-mer frequencies, and GC and AT-skew ^30–32^. These attributes are used as features in the supervised approaches, enabling the algorithms to learn virus signatures - patterns unique to viral genome sequences. Using class labels, e.g. phage and non-phage, of the sequences as inputs, these algorithms can learn these patterns and make predictions on novel sequences.

Virus identification methods can be broadly categorized into three groups based on their use of features: (1) reference-dependent methods rely on a reference database to generate features, (2) reference-independent methods generate nucleotide composition-based features without relying on a reference database, and (3) the hybrid methods rely on both reference-dependent and reference-independent features.

Reference-dependent methods distinguish viruses from non-viruses based on features identified in nucleotide or protein sequence databases. Such features might include similarity to known phages, hallmark phage genes, length of genes, and spacing between genes ^31,33,34^. Examples of reference-dependent tools are VirSorter2 ^31^, VIBRANT ^30^, and MARVEL ^33^. Although these methods report low false positive rates, they might miss novel or diverged viruses lacking similarities to viruses in the databases used ^29^.

In contrast, reference-independent or composition-based methods, such as VirFinder, while trained with a reference database in the training phase, do not rely on databases at run-time. Instead, using for example k-mer profiles or neural networks, these approaches generate numerical representations of query sequences, which are then used for classification ^32,35,36^. These methods are better at identifying divergent viruses than reference-dependent methods ^29^. Methods utilizing k-mer profiles as features also encounter several challenges. First, viral k-mer profiles may resemble those of their hosts^37–39^, making it difficult to distinguish viruses from their hosts based solely on k-mer frequencies. Next, although extending the length of k-mers can somewhat mitigate this problem, longer k-mers result in exponentially larger k-mer frequency vectors. This leads to sparse feature vectors, potentially causing machine learning models to become overfitted^40^. Lately, Convolutional Neural Networks (CNNs) have emerged for the de novo extraction of features to identify virus genome fragments, such as PPR-Meta, and DeepVirFinder ^35,36^. These CNNs can learn filters from training data, highlighting sequence motifs specific to viruses and other life domains. They condense this information into dense numerical vectors, addressing the limitations of k-mer profile-based machine learning approaches.

Finally, hybrid methods use reference-dependent and reference-independent features to identify phages. The recently published geNomad ^41^ approach belongs to this category. Such methods bring together the advantages of both worlds, and can be more sensitive than reference-dependent methods, and more precise than composition-based methods.

Several recent benchmarking studies compared the performance of state-of-the-art bioinformatic virus identification tools using simulated viral and non-viral testing datasets sampled from complete viral and microbial genomes, e.g. in the NCBI RefSeq database ^42–45^. These studies found that composition-based machine-learning methods outperformed reference-dependent methods. However, simulated data sourced from public databases presented several limitations: (1) Given that numerous virus identification tools utilize sequences from RefSeq for training, overlaps between testing datasets and training/reference datasets may introduce biases into the results, leading to more true positives. (2) The complexity of simulated sequence data does not match real-world metagenomic data. Wu *et al.* used real-world metagenomic and metaviromic benchmarking data from three biomes to avoid the above-mentioned biases. The authors noted that composition-based CNN exhibits greater TPR than other composition-based and reference-dependent methods. However, this heightened TPR comes at the cost of compromised precision. This might be due to the incompleteness of the training datasets and the lack of reliability checks. For instance, PPR-Meta did not include eukaryotic genomes in training datasets, which might lead to misclassification of eukaryotic sequences as viral sequences.

In this work, we present *Jaeger* (Just AnothEr phaGE findeR), a new tool utilizing a CNN to identify phage sequences among metagenome-assembled contigs. *Jaeger* ensures high TPR and precision by employing a dilated CNN with six-frame amino acid parameter sharing and integrating a reliability module. Besides, *Jaeger* incorporates ten times more parameters than existing CNN tools to capture the nuanced differences between viral and non-viral sequences. *Jaeger’s* training datasets encompass eukaryotic genomes, mitigating the misclassification of eukaryotic sequences as viral. *Jaeger* is the first deep-learning approach trained to directly recognize protein-level signatures from six-frame translated nucleotide sequences to identify viruses. *Jaeger* has been written in Python, is fully open-source, and is easy to install and use. Using GPU computing, Jaeger runs a lot faster than existing tools. Notably, Jaeger’s performance aligns with state-of-the-art approaches like PPR-Meta, DeepVirFinder, Virsorter2, and geNomad, while demonstrating a speed advantage approximately 20 times faster in CPU mode and up to 140 times faster with GPU acceleration. This allows Jaeger to be applied to vast datasets, as shown below.

## Results

### Distinguishing phages from cellular sequences with deep learning

Jaeger is a composition-based method for identifying contigs originating from phages in metagenome assemblies. This is achieved by a novel modular neural network architecture designed to capture protein-level features directly from nucleotide sequences (Figures 1, 2, and Supplementary Methods). The deep neural network (DNN) learns to identify amino acid motifs unique to each taxonomy category based on a training database of bacterial, archaeal, eukaryotic, and phage sequences. Our benchmarks show that this method outperforms traditional architectures that scan for nucleotide motifs (Supplementary Methods). We suggest that this reflects the higher conservation of protein than DNA sequences.

**Figure 1:**
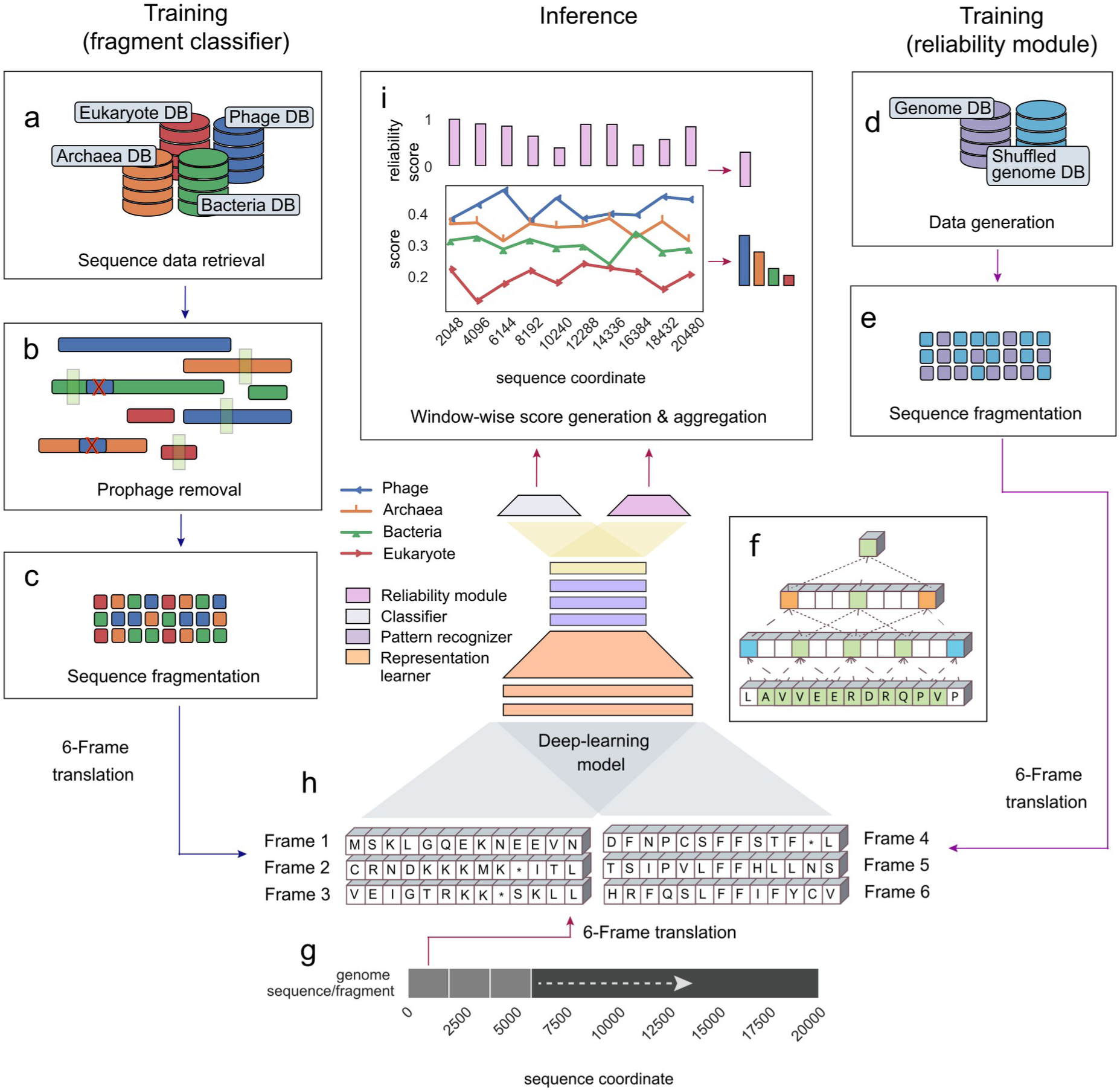
The Jaeger deep learning framework. The blue and magenta arrows show how training data is processed for nucleotide fragment classifier and reliability module training. The pink arrows show how an input nucleotide sequence is processed during inference. (**a**) Bacterial, archaeal, eukaryotic, and phage DNA sequences extracted from RefSeq and INPHARED ^46,47^ are used for training the model (see Methods). (**b**) Prophages in the genomes of bacteria and archaea are predicted ^31,48^ and masked ^48^. (**c**) All genomes are split into non-overlapping, 2,048 bp long fragments to create a nucleotide fragment database for training and testing. The panels on the right show the steps involved in the reliability predictor development. (**d**) Genome DB and Shuffled Genome DB are used to train the reliability predictor. Genome DB contains sequences from Phage DB, Eukaryote DB, Bacteria DB, and Archaea DB training sets. Shuffled Genome DB was created by randomly shuffling the sequences in the Genome DB. (**e**) All sequences are fragmented into non-overlapping 2,048 bp long fragments. A logistic regression model was trained using the feature vectors obtained for fragmented Genome DB and shuffled Genome DB from the fragment classifier. (**f**) Information propagation through dilated convolutional layers. Dotted lines link inputs to convolutional filters and their outputs. Dotted lines between the bottom layer and the one above depict standard convolution. Convolution with dilation rates of 2 (filter length=4) and 3 (filter length=3) are shown by dotted lines. (**g**) 2,048 bp long nucleotide fragments are extracted from the query sequence. (**h**) Each nucleotide fragment is translated into six reading frames and mapped to a learnable numerical vector. The neural network consists of a representation learner, a pattern recognizer, a logistic classifier, and a reliability estimator. The representation learner converts the six-frame translations to numerical vectors (amino acid embedding) as input to the pattern recognizer. The pattern recognizer recognizes hidden patterns in the sequences and converts that information into a numerical vector (sequence embedding). The logistic classifier utilizes the sequence embeddings to make predictions. ( **i**) The reliability module estimates the likelihood of the projections made by the logistic classifier being a true positive. The model outputs five floating point numbers per non-overlapping 2,048 bp fragment by default, representing phage, bacterial, eukaryotic, archaeal, and reliability scores, respectively. The taxonomy class with the maximum mean score is chosen as the putative taxonomy class for the input sequence.

**Figure 2.**
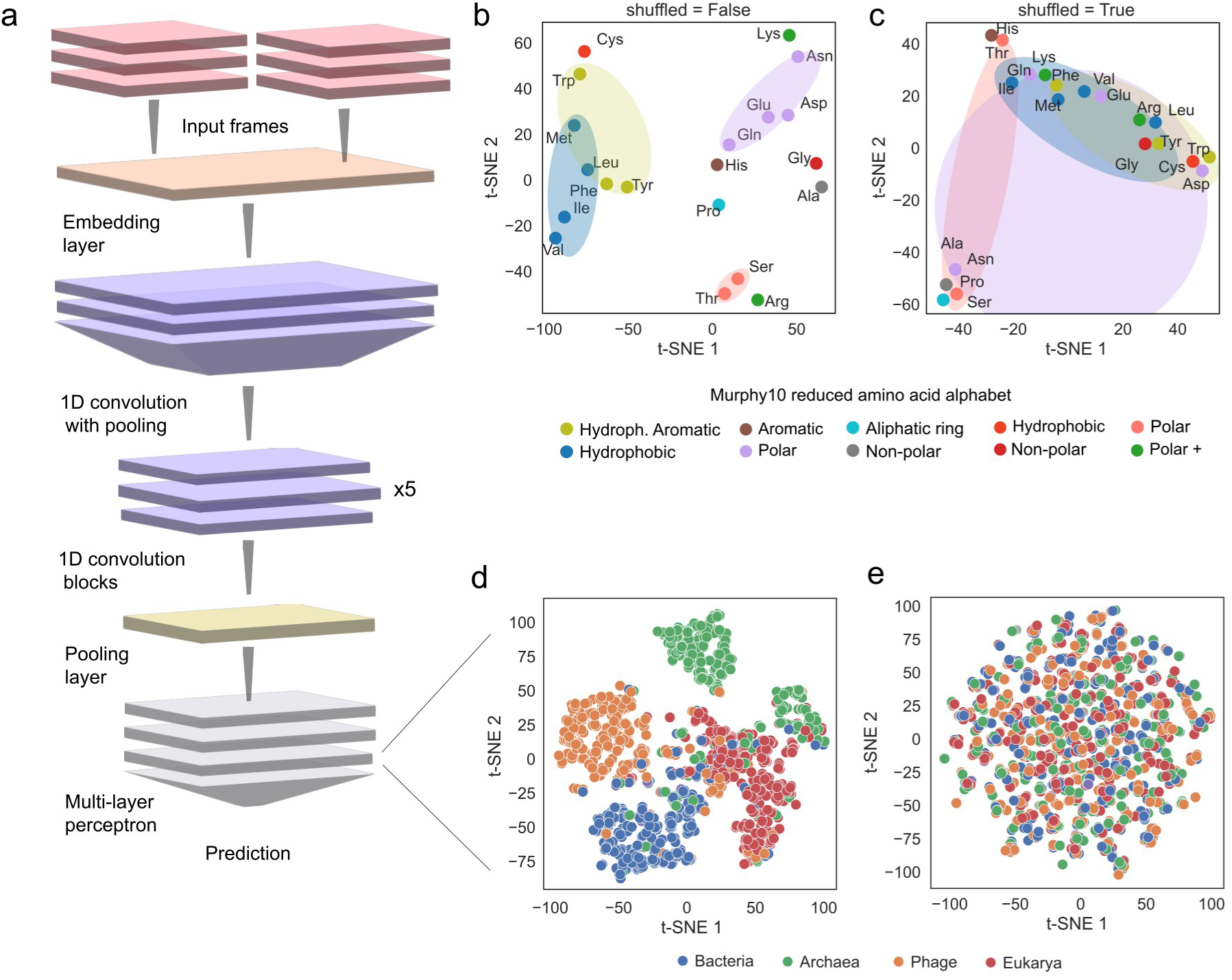
**(a**) Representation of the different DNN sections implemented in Jaeger. See Methods and Supplement for details. (**b**) The T-SNE projection of Jaeger’s amino acid embedding vectors reveals a distinct pattern where amino acids sharing similar physicochemical properties are closely grouped in the embedding space. The coloring of amino acids follows the Murphy10 reduced amino acid alphabet (Murphy et al., 2000). (**c**) When the embedding vectors were randomly permuted, clustering among similar amino acids was lost. (**d**) The T-SNE projection of sequence embedding vectors extracted from the neural network during inference shows the clustering of sequences from the same category. (**e**) When the embedding vectors were randomly permuted, the clusters in the T-SNE projection collapsed.

Machine learning tools should inform users whether they are uncertain about a prediction. This avoids over-interpreting weak, potentially spurious predictions. To achieve this, we have developed a method (see Methods) to measure the reliability of the neural network’s predictions. This metric - called the reliability score - alerts users when the network is not entirely confident in its assessment. In such cases, human intervention may be required to confirm the predictions.

Jaeger’s pipeline consists of three steps. First, preprocessing of genomic sequences; second, sequence classification and reliability estimation; third, aggregation of scores and output generation (Figure 1). In the preprocessing stage, all input sequences longer than the window length (2,048bp) are split into non-overlapping fragments. The neural network assigns four scores to each fragment, indicating the likelihood of belonging to bacteria, archaea, eukaryotes, and phages. Next, the reliability predictor is used to quantify the confidence in neural network-predicted scores. The final prediction and the reliability score for a contig are calculated by taking the mean of all fragment scores per contig. Then the sequence is assigned to the class with the highest mean predictive score. Jaeger generates an output table indicating the final prediction for each contig, average prediction scores, and average reliability scores. Users can request window-wise prediction scores if needed.

### Estimating the reliability of neural network predictions

It has been observed that neural networks may display high confidence even when they make incorrect predictions, especially when the distribution of new inputs is significantly different from that of the training data. Given the vastness of the biological sequence space and the limited subspace on which the network was trained, highly dissimilar sequences are likely in real-world applications. Several methods have been proposed to identify when a neural network is uncertain about a prediction ^49–51^. For our classification task, we used the Neural Mean Discrepancy method ^52^, which analyzes the output values of the neural network’s penultimate layer for differences between in-distribution and out-of-distribution samples (Supplementary Figure S1). The reliability module (Figure 1i) outputs a number between 0 (least confident) and 1(most confident) that quantifies the neural network’s uncertainty. This helps users to assess Jaeger’s predictions and assess their reliability.

### Interpreting neural network’s predictions

Deep learning models are often viewed as black boxes; however, significant efforts have been made to enhance their interpretability. One such method is gradient-based interpretation, also referred to as saliency mapping ^53^. This approach involves analyzing the gradients of a model’s output with respect to its input features to understand the underlying prediction mechanisms. By calculating these gradients, we can identify the features that most influence the model’s decision. We observed a moderate correlation (r=0.49) between gradient magnitude and gene density. Suggesting that the neural network allocates attention to each genomic strand in proportion to its gene density (Supplementary Figure S2, Supplementary Methods). Therefore, it incorporates protein motif information from both strands to make predictions.

### Benchmarking Jaeger on simulated metagenome assemblies

Microscopic eukaryotes are ubiquitous in biomes such as seawater and soil ^54,55^. However, many existing composition-based tools are trained only with prokaryotic and viral sequences. This could result in incorrect identification of eukaryotic sequences as having a phage origin. To address this problem, we trained Jaeger with additional eukaryotic genomes. To compare Jaeger’s and other methods’ performance in classifying sequences from different domains, we simulated ten replicate contig datasets from three biomes by selecting and fragmenting genomes of prokaryotes, eukaryotes, and phages (see Methods, Supplementary Data 1). For sequences from every domain (Figure 3a) and every simulated biome (Figure 3b), Jaeger performed better than state-of-the-art composition-based tools (Seeker, PPR-Meta, and DeepVirFinder) and similarly to reference-dependent methods (VirSorter2 and geNomad). All methods except Jaeger and Virsorter2 misclassified many eukaryotic sequences as being of phage origin (Figure 3a). Thus, Jaeger avoids over-predicting eukaryotic sequences, reducing the risk of identifying microscopic eukaryotes falsely as phages. Jaeger had the lowest proportion of misclassifications in both the simulated soil and seawater metagenomes, whereas VirSorter2 showed the best performance for the simulated human gut dataset, where Jaeger predicts that approximately 15% of the prokaryotic contigs are viral, whereas VirSorter2 mispredicts only 3%. This difference in performance might be due to the specifics and small size of the simulated dataset used for benchmarking (150 prokaryotic, 6 eukaryotic, and 150 phages, see Methods). Below, we show additional results of a second, less biased benchmark using real-world metagenomic sequences.

**Figure 3.**
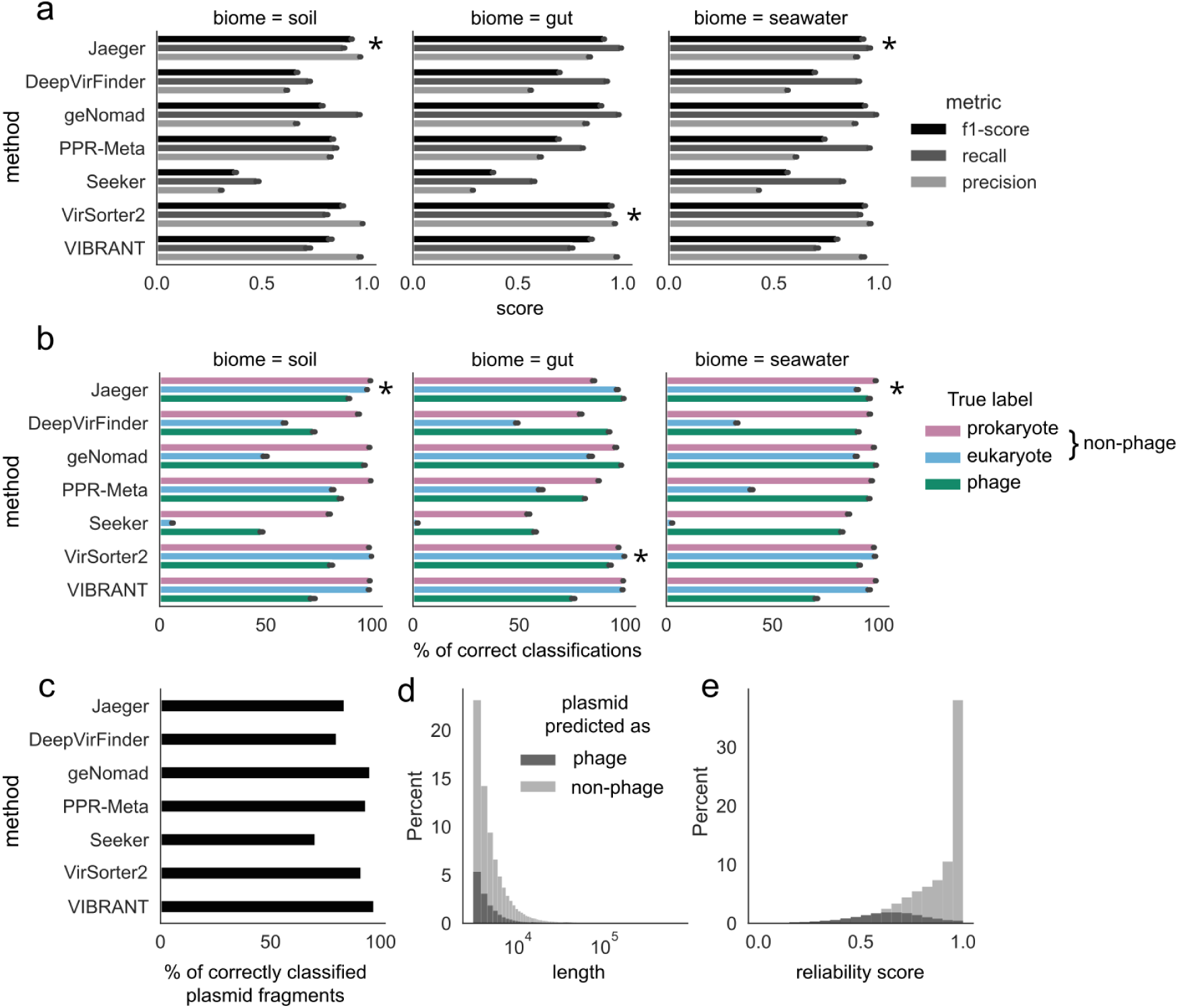
Genomic sequences of bacteria (n=150), eukaryotes (n=6), and phages (n=150) with biome annotations were selected and fragmented to simulate metagenomic contigs. (**a**) Performance metrics (average of 10 replicates) of different methods on the simulated metagenomic datasets. * denotes the method with the best F1 score for the respective category. (**b**) Domain level accuracy of each method on the simulated metagenomic datasets (average of 10 replicates). The green columns refer to the percentage of correctly classified phage contigs. The blue and purple columns show the percentage of eukaryotic and prokaryotic contigs correctly identified as non-phage. Jaeger and Virsorter2 misclassified fewer eukaryotic contigs as viral origin than the other methods (**c**) Plasmid sequences from RefSeq were fragmented to simulate plasmid contigs. The effectiveness of each method in distinguishing plasmids from phages was then evaluated. The percentage of plasmid contigs that each method correctly classified as non-viral is presented here. (**d**) Length distributions of plasmid fragments classified as phage and cellular by Jaeger. (**e**) Reliability score distributions for plasmid fragments classified as phage and cellular by Jaeger. Higher reliability scores indicate high confidence in predictions.

### Benchmarking Jaeger’s sensitivity to sequencing errors

To assess Jaeger’s sensitivity to sequencing errors, we artificially mutated the simulated metagenome-assembled contig datasets at mutation rates ranging from 0.001 to 0.2 *in silico* by randomly replacing the original nucleotide with another. Results (Supplementary Figure S3) show that at mutation rates below 0.05, the F1 score drop is between 0.002% and 1.5%. At a mutation rate of 0.05, the F1 score drop is within 0.17 to 6%. At 0.1 mutation rate, the drop in the F1 score is between 6.17 to 12.95%. The model’s performance drops sharply at a mutation rate of 0.2, evidenced by the 27.66 to 40.87% drop in the F1 score. Error rates of commonly used sequencing platforms lie within the 0.001 to 0.1 range. The results indicate that Jaeger’s predictive capability remains consistent within the range of error rates of widely used sequencing platforms.

### Benchmarking Jaeger on the IMG/VR virus database

As many virus identification tools included the RefSeq data in their training/reference databases, using RefSeq sequences to test them might bias the results (more true positives expected). Besides, the RefSeq database contains genome sequences from many cultivated and isolated viruses, while those in environmental samples remain under-characterized. To avoid biases in our comparison, we further benchmarked tools using viruses from IMG/VR ^56^.

IMG/VR is a large database of prokaryotic and eukaryotic viral genomic sequences, predominantly identified in metagenomics data. The database also provides the quality tier of the sequences based on completeness estimated by CheckV ^57^. Since Jaeger is only trained to detect phages, we selected all the genome sequences (n=178,139) of phages categorized by CheckV as high-quality (>90% complete) excluding viruses also present in the RefSeq dataset for benchmarking, hereafter called the virus database. A large fraction of this dataset were prophage sequences (n=78,518). Here, we tested geNomad’s neural network classifier (geNomad-nn) and geNomad’s hybrid classifier (neural network classifier + marker-based classifier) separately.

GeNomad (hybrid) recorded the best recall (0.99) over almost every length range, followed by Jaeger (0.94) (Figure 4a). This high performance was expected since most of the sequences in the IMG/VR database were predicted using geNomad ^56^. The recall of all methods was not related to the genome length buckets. This is likely because some methods are less sensitive to different taxonomic groups of phages, as the genome lengths of taxonomically related phages tend to fall within the same range.

**Figure 4.**
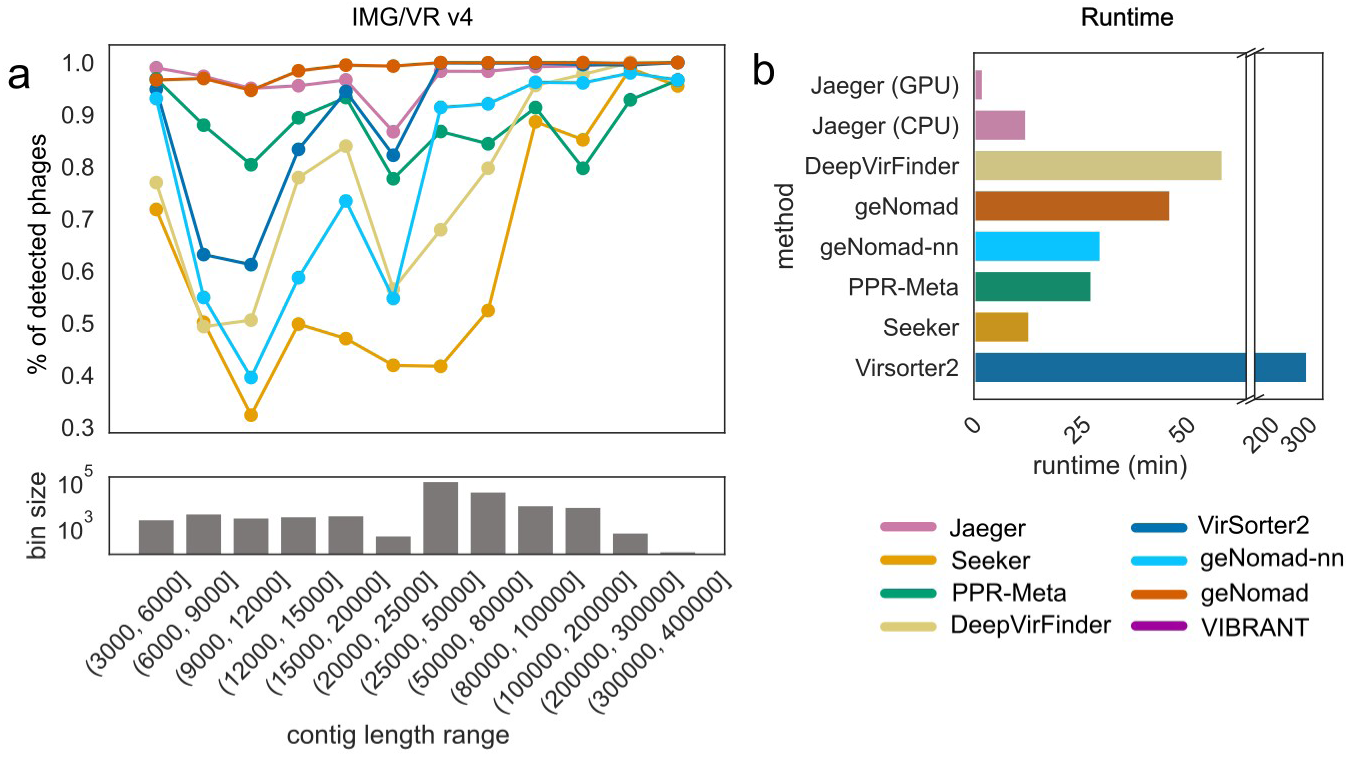
Jaeger’s performance was compared to five other state-of-the-art methods on the prokaryotic virus sequences that CheckV annotated as “high quality” (n=178,139) from the IMG/VR (v4) database. (a) The ability of different methods to correctly identify all “high-quality” viral sequences at different length buckets. (b) Run time was measured by wall clock time in seconds. Jaeger recorded the shortest runtimes in both GPU and CPU modes.

Jaeger exhibited the best performance-to-speed ratio among all the methods we benchmarked. In CPU mode, Jaeger was more than 3.5 times faster than geNomad and more than 20 times faster than Virsorter2, while with GPU acceleration, Jaeger surpassed geNomad by over 19 times and Virsorter2 by over 140 times in terms of speed. Jaeger’s high speed and accuracy position it as an attractive choice for large-scale metagenomic virus discovery and surveillance projects.

### Benchmarking Jaeger on real-world metagenomic data

As both the macro- and micro-diversity of viruses in natural samples are usually much greater than the diversity of viruses in the database ^58^, this further obscures the benchmarking results. To avoid biases in our comparison and reach the complexity level of microbial/viral communities in real metagenomic datasets, we utilized three real-world datasets from soil, gut, and seawater biomes. Each biome dataset comprised eight viral and microbial dataset pairs, complementing the data from simulated metagenomes and the virus database. The details of these three datasets can be found in Wu *et al* ^29^. Consistent with this previous benchmark, gut samples are poorly predicted by all tools. This could be due to the contamination of microbial elements in the gut viral datasets^29,57,59–62^, which contained the lowest numbers of viral markers and the highest numbers of microbial markers among the three biomes^29,57,59–62^. Jaeger detected many contigs from the viral size fraction and few from the microbial size fraction (Figure 5), indicating high TPR (TP/(TP+FN)) and high specificity (TN/(TN+FP)), respectively^29^. In one of the seawater samples, Jaeger detected 96% of viruses from the viral fraction (TPR) and had a relatively high precision (TP/(TP+FP)) of 81% (Supplementary Table 1). Jaeger performed consistently well on soil samples, with 89% ± 4% of viruses detected from viral fractions and 1.57% ± 0.60% of viruses detected from the microbial fractions. Jaeger achieved high TPR (seawater: 88% ± 4%, soil: 86% ± 0.5%), specificity (seawater: 95% ± 1%, soil: 99% ± 0.5%), precision (seawater: 85% ± 7%, soil: 99% ± 0.5%), and F1 score (seawater: 87% ± 5%, soil: 92% ± 0.4%) on both seawater and soil biomes (Supplementary Table 2). Overall, we estimate a false discovery rate (FP/(FP+TP)) of 8%, although we note that these numbers strongly depend on the biome and the specific studies (see Table 1 and Table 2). Jaeger outperformed all the other state-of-the-art tools in F1 score, including the recently published tool geNomad. VirSorter2 showed the lowest FDR (0.02) but had lower sensitivity than Jaeger. Notably, Jaeger achieved this high performance on real-world datasets, even though the training data consisted solely of genome sequences from the RefSeq and INPHARED databases. This highlights the potential of our neural network to extract universal phage features from the sequence data.

**Figure 5.**
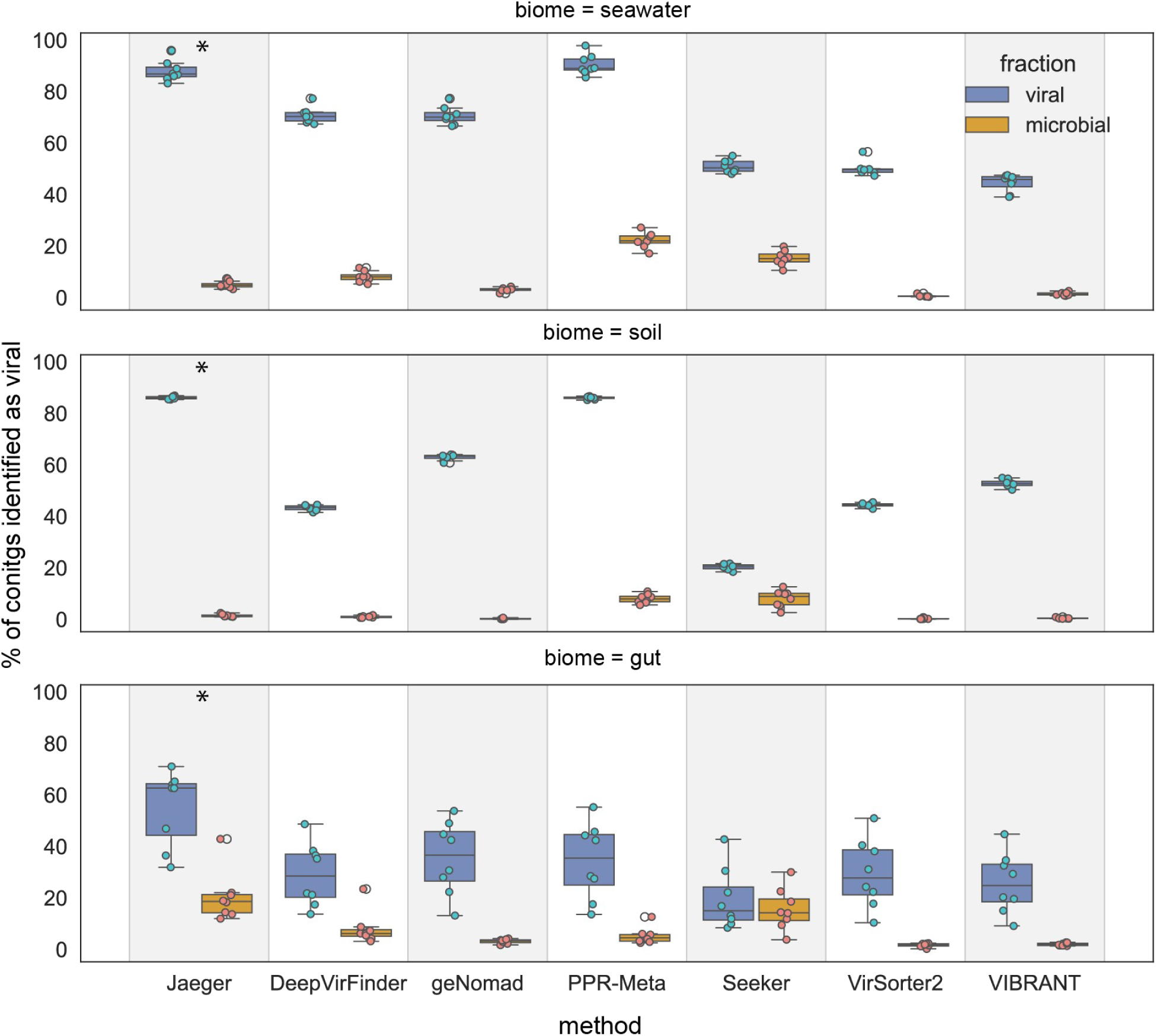
Percentage of contigs identified as viral in the viral (true positive rate, blue) and microbial (false positive rate, orange) datasets in three biomes. * denotes the method with the biggest difference between the TPR and FPR. For details on the approach, see ref ^29^.

**Table 1.**
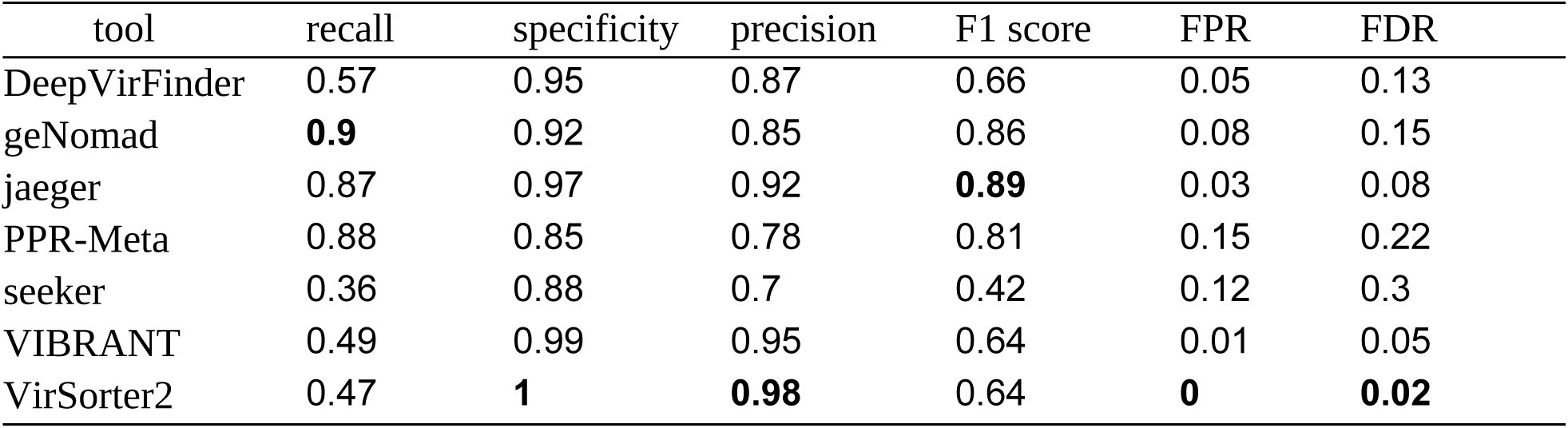
Mean metrics of real-world metagenome benchmark (seawater and soil biomes). For details see Supplementary Table 4.

| tool | recall | specificity | precision | F1 score | FPR | FDR |
| --- | --- | --- | --- | --- | --- | --- |
| DeepVirFinder | 0.57 | 0.95 | 0.87 | 0.66 | 0.05 | 0.13 |
| geNomad | **0.9** | 0.92 | 0.85 | 0.86 | 0.08 | 0.15 |
| jaeger | 0.87 | 0.97 | 0.92 | **0.89** | 0.03 | 0.08 |
| PPR-Meta | 0.88 | 0.85 | 0.78 | 0.81 | 0.15 | 0.22 |
| seeker | 0.36 | 0.88 | 0.7 | 0.42 | 0.12 | 0.3 |
| VIBRANT | 0.49 | 0.99 | 0.95 | 0.64 | 0.01 | 0.05 |
| VirSorter2 | 0.47 | **1** | **0.98** | 0.64 | **0** | **0.02** |

### Benchmarking Jaeger on plasmid fragments

Plasmids are often considered different from phages but can have similar features, making it difficult to distinguish them ^63^. Furthermore, hybrids with phage-like and plasmid-like characteristics, called phage-plasmids exist in nature ^64,65^, making the distinction even more challenging. We tested the ability of various methods to identify a non-redundant set of plasmid sequences downloaded from RefSeq (n=55,417). We created a plasmid fragment dataset with the same empirical length distribution as the simulated metagenomic contigs (n=624,904, median length=4,614, Q1=3,585, Q3=7,157, max=695,420). Here, we measured how often tools misclassified plasmid fragments as having phage origin. As the classification rates for TP and FP are related and should both be considered when interpreting false predictions, we also mention the mean TPR and FPR values that were determined by benchmarking on real-world metagenomic data above. Figure 3c shows that VIBRANT with virome mode misclassified the lowest percentage, but also had a low TPR (2.75%, TPR=0.41, FPR=0.01), followed by geNomad (4.5%, TPR=0.56, FPR=0.02), PPR-Meta (6.4%, TPR=0.7, FPR=0.12), VirSorter2 (8.5%, TPR=0.41, FPR=0.01), Jaeger (16%, TPR=0.76, FPR=0.12), DeepVirFinder (19%, TPR=0.48, FPR=0.06), and Seeker (29%, TPR=0.3, FPR=0.13). Based on these results, the percentage of falsely classified plasmids correlated positively with the previously determined FPR (R^2^=0.41) while the correlation with TPR was slightly negative (R^2^=0.10), suggesting that methods with low FPR misclassified fewer plasmids as phages, but this did not strongly impact TPR. While other methods processed all 624,904 fragments in the plasmid dataset, VIBRANT only processed 108,533 fragments, so its effective percentage of misclassifications was 15.8%. This difference is likely because VIBRANT requires at least four open reading frames to be present on a contig to make a prediction. As expected, methods designed to identify both plasmids and viral sequences, such as geNomad and PPR-Meta, performed the best in this benchmark. All methods misclassified shorter fragments more often than longer ones (Figure 3d). For instance, plasmid fragments misclassified as prophages by Jaeger had a median length of 4,279 (Q1=3,484, Q3=6,141), while the median length of correctly classified fragments was 4,685 (Q1=3,610, Q3=7,380). This is probably due to the lack of plasmid-specific signals in short contigs. Surprisingly, Virsorter2 also reported few plasmid misclassifications, even though not trained on plasmid sequences. While Jaeger outperformed other composition-based methods like DeepVirFinder and Seeker, it still had quite a high misclassification rate. Although plasmids are included in their respective host domain datasets, our neural network did not include a separate plasmid training class, so it may not have learned enough features to distinguish them from phages. Figure 3e shows that the reliability module assigns lower reliability scores to plasmids predicted as phage sequences. Based on this analysis, we set the default reliability score cutoff at 0.2.

### Global metagenome assemblies contain 11.3% phage contigs

The GPU mode and the resulting high speed enable Jaeger to predict phage sequences in massive datasets. To test the scalability of Jaeger, we set out to identify phages in all 16,702 publicly available assembled metagenomes from the MGnify database as of August 2023 (Figure 6, Supplementary Table 3, ^66^).

**Figure 6.**
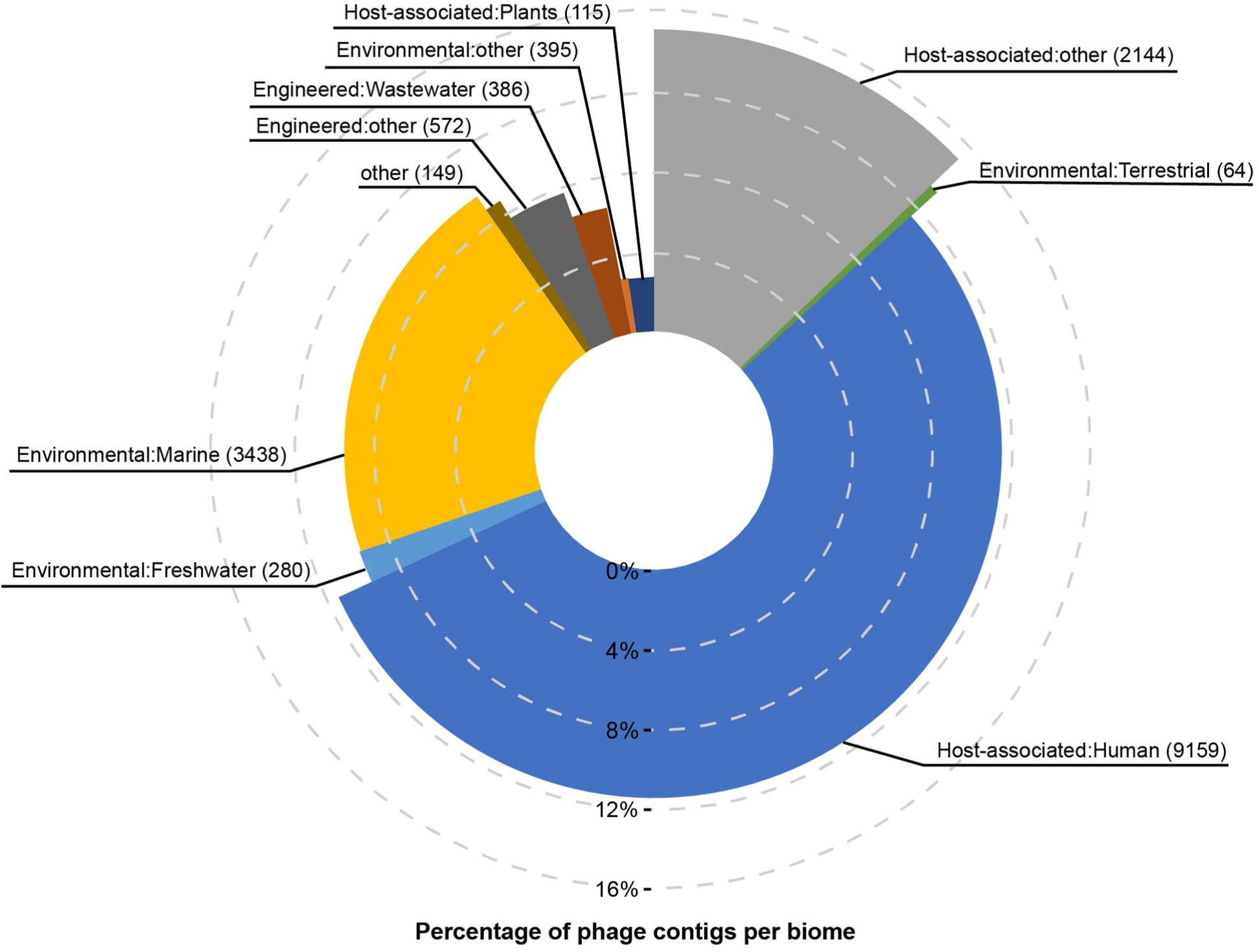
Results of running Jaeger on 16,702 publicly available assemblies in the MGnify database. The angle of each slice indicates the number of datasets per biome (number between brackets) and the radius indicates the average percentage of predicted phage contigs.

The MGnify assemblies contained predominantly short sequences. In total, >143 million contigs were longer than 2,048 bp and 875 were longer than 1Mb (Supplementary Table 3), with a median contig length of 3,448bp across all samples. Within 394 GPU hours, Jaeger predicted phage sequences among the MGnify contigs. The runs used a maximum of 9.3 GB RAM. Remarkably, 11.3% of all contigs were predicted as phage. The fraction of phage sequences ranged from 0-100% per dataset, with most assemblies containing 10^3^ to 10^5^ contigs ≥2,048bp. Most public metagenomes were host-associated (11,418 or 68%, Figure 6). Human metagenomes contained on average 11.5% phage sequences, plant-associated biomes 2.7%, and other hosts 15.2% (Figure 6). Note that the abundance of the contigs was not taken into account. Different contigs may have different vertical coverage or read depth, which reflects sequence abundance in the metagenome. Thus, the radius of the slices in Figure 6 should not be confused with the overall fraction of viral DNA. Moreover, fragmented genomes, such as those derived from diverse populations, may contribute more contigs than contiguous genomes, such as those derived from uniform populations. Together, we conclude that the average fraction of predicted phage contigs represents each biome’s (recognized) viral diversity.

It is important to note the variation among biomes in the average percentage of phage contigs. Host-associated metagenomes, particularly of mammal origin, have more phage contigs than metagenomes from most other biomes. We hypothesize that this reflects (1) ecological differences in the abundance and diversity of bacteriophages, and (2) the fact that phages in these biomes are traditionally better studied, and thus better represented in the database. Despite our efforts to reduce redundancy in the training datasets (see Methods), the latter may still be expected to play a role. As more metagenomes from diverse environments are published and methods to identify phages improve, the percentage of phage sequences in different biomes will likely be refined. Additionally, with the advances in accurate long-read sequencing and ongoing improvements in bioinformatic sequence assembly tools, the contigs obtained from publicly available metagenomes will become more representative of their respective microbiomes and viromes. This will improve estimates of the prevalence of phages across different environments.

## Discussion

Bacteriophages play vital roles in nearly every biome on Earth by interacting with microbial communities. Different tools have been introduced to identify phages in metagenomic datasets, and they all have their strengths and weaknesses. Our recent benchmarking study showed that deep learning methods are more sensitive than reference-dependent methods at the cost of specificity ^29^. This could be addressed by (1) improving the neural network architecture, (2) including microbial eukaryotes (fungi and protists) in the training dataset, and (3) implementing measures to improve the reliability of the predictions. These features were included in our newly presented tool Jaeger.

We compared Jaeger to other state-of-the-art methods using a range of datasets and benchmarking methods and showed it has better or comparable performance. Jaeger outperformed all tested deep learning viral prediction tools, showing a 1.5-7% improvement in F1 score for real-world metagenome datasets. Jaeger’s performance is comparable to state-of-the-art homology-assisted methods while being up to ∼20x faster in CPU mode and ∼140x faster with GPU acceleration on a laptop computer. While the 8% false-discovery rate means that not each predicted sequence should be taken for granted, its high speed and accuracy make Jaeger a promising tool for virus discovery by metagenomics, especially in high-throughput virus discovery efforts with massive datasets.

## Methods

### Datasets for training, validation, and benchmarking

Genome assemblies of Bacteria, Archaea, Fungi, and protists from the SAR supergroup were retrieved from RefSeq (March 2022). Fungal and protist genomes were combined to create the Eukaryote dataset. We picked contigs longer than 20,000 bp from fragmented assemblies to minimize noise. We randomly sampled one genome per genus to reduce redundancy. This resulted in a dataset of 1,559 (mean length=4,072,688 bp) bacterial, 128 (mean length=2,823,770 bp) archaeal, and 117 (mean length=37,828,892 bp) eukaryotic genome sequences (Supplementary Data 2). RepeatMasker was used to mask low-complexity repeat regions of Eukaryote genomes. Prophages in prokaryotic genomes were identified using PHASTER and Virsorter2 (v2.2.3) and were masked with N characters using a custom script. Genomes of phages (n=14,041, mean length=60,764 bp) were retrieved from the INPHARED (v 1.7) database. The redundancy of the phage sequences was removed using dedupe from BBTools (v38) at a minimum identity of 95%. As we aimed to train a genome fragment classifier, we created artificial fragment databases for model training and evaluation. All genomes were fragmented into non-overlapping short sequence segments of 2,048 bp to and clustered using MMseqs2 (release 13, mmseqs2 easy-cluster -s 2 –min-seq-id 0.9 -c 0.8) to remove similar sequence fragments. This resulted in four fragment databases: Phage DB (n=265,959), Eukaryote DB (n=1,375,939), Bacteria DB (n=915,817), and Archaea DB (n=459,660). Each non-redundant fragment database was split into three subsets to be used for training (70%), validation (5%), and testing (25%). Random subsets from each taxonomic group were combined to create the final datasets.

Real-world data for benchmarking were obtained as described before ^29^. Briefly, paired real metagenomic datasets were derived from recent virome studies on seawater ^67^, soil ^68^, and gut ^69^, three highly divergent biomes ^70^. Raw reads were pre-processed and assembled into contigs. Only contigs with lengths of at least 2,048 bp were used. Homologous contigs between viral and microbial fractions were removed ^29^. Simulated metagenome-assembled contig datasets representing three biomes were created as follows.

Prokaryotic (n=70), eukaryotic (n=6), and phage (n=150) genomes and their associated biome information were obtained from GTDB, RefSeq, and IMG/VR, respectively (Supplementary Data 1). These were fragmented into scaffolds according to an empirical contig size distribution from the real metagenomic assemblies (min length=3,000, mean length=13,402, max length=729,405, bins=100,000). For each biome, up to 10,000 scaffolds were sampled in ten replicates to account for the variability in the sampling process. Metagenome assemblies were downloaded from MGnify ^66^ using a custom script through the MGnify API. High-confidence prokaryotic viral contigs were obtained from the IMG/VR (V4) database.

### Training deep learning models

The data preprocessing pipeline was implemented using custom Python scripts. The sequence data loader was implemented by subclassing the TensorFlow Data class. The data loader translates nucleotide sequences along six reading frames and maps each amino acid to an integer value. An Embedding layer was used to map the integer representation of each six-frame translated amino acid vector to a learnable embedding (LE) vector consisting of 4 floats. Thus, input vectors to amino acid-based models have three dimensions (n_frames, n_codons_in_sequence, dim_LE_vector). Input vectors to nucleotide-based models have three dimensions (strand, sequence_length, nucleotide_representation). For a sequence of 1,024 bp, these vectors would have dimensions of (6, 340, 4) and (2, 1024, 4) respectively.

All models were implemented in TensorFlow 2.10 ^71^ and Keras 2.10 ^72^ and trained using a supervised learning approach. The Dilated Convolutional Neural Network with Residual connections (DCNN-Res) used here (Figure 2) contained 947,040 (trainable=943,964, non-trainable=3,076) parameters distributed over 28 layers. A complete description of this model’s topology can be found in **Supplementary Methods**. Initial weights for models were initialized by sampling from a He-Uniform Distribution. To prevent overfitting, ridge regularization was applied to kernel weights (l2=1e-5) and bias weights (l2=1e-2). Dropout regularization of rate 0.1 is applied after the max pooling layer, and after the fully connected layer, to reduce overfitting in neural networks by randomly dropping out a few dimensions. Since the training dataset is imbalanced, the bias of the output layer is set to reflect this imbalance. The model was trained by minimizing the cross-entropy loss between the predicted score and the true labels (0 for Bacteria, 1 for Phage, 2 for Eukaryotes, and 3 for Archaea). The training dataset is iteratively fed into the model in batches of size 32. The parameters in the neural networks were updated through back-propagation using the Adam optimization algorithm for stochastic gradient descent with ^73^ with a custom learning rate schedule. The learning rate was set to 5e-4 for the first 1000 optimization steps, and then we exponentially lowered the learning rate to 1e-4. Hyperparameters for all the models were selected by evaluating the performance against the validation set. Early stopping criteria were used to stop training once the model performance stopped improving on the validation set to prevent overfitting. Please refer to the Supplementary Methods section for additional details on the model architecture.

### Ensuring the reliability of predictions with a Reliability Model (RM)

We implemented a post-hoc, auxiliary multiple logistic regression model **Eq. 1** using neural activation statistics to calculate a reliability score for every prediction ^52^.

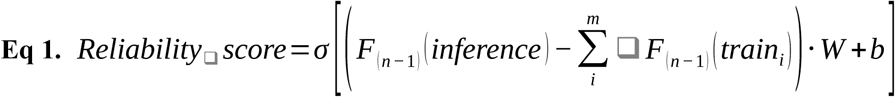

where,

- *F*_(*n−*1)_ is the output of the penultimate layer of the neural network,
- *σ* is the sigmoid function,
- *inference* refers to the sequence for which the reliability score needs to be calculated,
- *train_i_* is the *_i_^th^* sequence in the neural network’s training dataset,
- *m* is the total number sequences in the training dataset,
- *W* represents the weights of the reliability module,
- *b* b represents the biases of the reliability module.

The reliability score warns users of less reliable predictions, reducing the risk of incorporating erroneous sequences into downstream analysis. This model will be hereafter referred to as the reliability model (RM). The positive dataset to create the reliability estimation model was generated by randomly sampling 500,000 sequence fragments from training datasets. The negative dataset (n=500,000) was generated by randomly permuting the positive sequences and generating random sequences without varying the G+C content using a custom script. Embedding vectors (dim=128) for each fragment in the resulting datasets were obtained by the penultimate layer of the neural network. Next, embedding vectors were l2 normalized ^74^. Next, the positive and negative datasets are combined and split into training (70%) and testing (30%) datasets.

Generated features were used to train a logistic regression model with the Scikit-learn 1.3.2 library ^75^. Hyperparameter search was performed using 5-fold cross-validation (optimum parameters; C=25, solver=’lbfgs, penalty= ’l2’). Platt’s method, as implemented in Scikit-learn via the ’CalibratedClassifierCV’, was utilized to calibrate the probabilities of the final model ^76^.

### Jaeger uses a sliding window approach to give final predictions

Jaeger employs a non-overlapping sliding window to generate scores across a query sequence. By default, the length between two consecutive windows is 2,048bp. Users can change this using the –stride flag. For every 2048bp window, the neural network returns four scores, corresponding to its trained classes (Bacteria, Archaea, Eukaryote, and Phage). Specifically, we split long sequences into shorter fragments of the size of the input layer using a sliding window (2048bp). Each short fragment is then processed by the DLM to obtain a predictive score and feature embeddings. To calculate the reliability score, we utilize feature embeddings. We then aggregate the predictive scores and reliability scores of all windows to calculate the mean predictive and reliability scores. Finally, we assign the sequence to the class with the highest mean score (Figure 1). We note that advanced users may use the per-window scores to assess potentially integrated prophages in longer host contigs. These can be accessed using the –getalllogits flag.

### Performance benchmark

Jaeger (version 1.1.30a) was benchmarked against Virsorter2 (version 2.2.4), PPR-Meta (version 1.1), DeepVirFInder (version 1.0), and Seeker (version 1.0.3), and geNomad (version 1.7) ^31,35,36,41,77^. Default cutoffs were used for tools that output a prediction score. Any contigs that were not included in Virsorter2’s output were regarded as being of cellular origin. Predictions from VIBRANT’s VIBRANT_machine_<filename> output were obtained, and we assumed that any contigs not included in this file were also of cellular origin. As for other tools, their default output files were parsed without making any assumptions. All outputs were aggregated, and performance matrices were calculated using custom scripts.

### Runtime comparison

For the runtime comparison of different tools, we sampled, 10,000 contigs from the high-confidence contigs of IMGV/R (v4) database (host=’bacteria’, average length=16,628 ± 20,130). All tools were run on a consumer laptop with an Intel i7 9750H processor, 32GB of RAM, and an NVIDIA 1660 Ti GPU with 6GB of memory.

### Global MGnify analysis

For the global analysis of metagenomes, every publicly available assembly in MGnify was downloaded iteratively through the API, resulting in 16,702 assemblies. We created a parallelized version of Jaeger which can utilize multiple GPUs. This parallelized version was run on a GPU node with four state-of-the-art A100 GPUs. Jaeger was then run on all the assemblies using default settings (Jaeger --stride 2048 --size 2048) and could predict 143,342,505 contigs with a minimum of 2,048bp length.

## Data availability

The data generated in this study have been deposited in the Zenodo database under doi: 10.5281/zenodo.13338131. The accession numbers of genomes used in this study can be found in Supplementary Data 2.

## Code availability

All the models and scripts developed in this study are available at https://github.com/MGXlab/Jaeger. The paper code release is also available at https://zenodo.org/records/13336195.

## Acknowledgments

The authors would like to thank the members of the Kaderali group, the Viral Ecology and Omics (VEO) Group at Friedrich Schiller University Jena, and the Theoretical Biology and Bioinformatics Group (TBB) at Utrecht University. Special thanks to Jankees van Amerongen (TBB) for invaluable support with big data and computer lab access. This project is supported by the European Union’s Horizon 2020 research and innovation program, under the Marie Skłodowska-Curie Actions Innovative Training Networks grant agreement no. 955974 (VIROINF), the European Unioin’s HORIZON-HLTH-2023-DISEASE program (APPEAL project, grant 101137311), the Utrecht University One Health Initiative, ZonMw project 541003001, the European Research Council (ERC) Consolidator grant 865694: DiversiPHI, the Deutsche Forschungsgemeinschaft (DFG, German Research Foundation) under Germany’s Excellence Strategy – EXC 2051 – Project-ID 390713860, and the Alexander von Humboldt Foundation in the context of an Alexander von Humboldt-Professorship founded by German Federal Ministry of Education and Research.

## Author contributions statement

Y.W. designed the analysis, carried out the data compilation, and performed the model construction. Y.W. and L.W. did the benchmarking, and testing, and wrote the manuscript. R.B. conducted a test and provided feedback. P.R. and Y.W. performed the synthetic benchmarks. S.D. generated the MGnify assembly dataset and calculated CPU/GPU usage. E.H. analyzed the Jaeger results on MGnify assemblies. L.K. and B.E.D. conceived the study and contributed to study design, data interpretation, and manuscript preparation. All authors read, edited, and approved the final manuscript.

## Notes

https://zenodo.org/records/13338131

### Competing Interest Statement

The authors have declared no competing interest.

