## Supplementary Methods for "Jaeger: an accurate and fast deep-learning tool to detect bacteriophage sequences"

### Determining the optimal neural network architecture for Jaeger

In this study, we compared four different neural network architectures - the baseline model with only a representation learner and a logistic regressor, the Dilated Convolutional Neural Network with Residual connections model (DCNN-Res), the DCNN-Res and LSTM model (DCNN-Res-LSTM), and the DCNN-Res and Attention model (DCNN-Res-Attention). Detailed configuration of each model is available in **Table S6**. Each architecture was tested with two different input representations of the nucleotide sequences for training the networks, the nucleotide- and amino acid-level representation. The embedding layer of each model converts nucleotides into a one-hot vector representation and amino acids into a randomly initialized learnable vector representation. The results showed (**Table S5**) that the DCNN-Res model is best suited for identifying viruses, possibly due to its ability to identify amino acid sequence motifs representing the different classes in our training dataset. Nevertheless, it is important to acknowledge that the lower performance of other architectures might also be attributed to the limitations of the optimization algorithm, which may encounter challenges in converging other architectures to lower minima.

After selecting the best model architecture, we extensively tested our final models with simulated metagenomic datasets, high-quality prokaryotic viruses from IMG/VR(v4) [1], three real-world metagenomic datasets [2], and 16,702 metagenomes from MGnify (see main text).

### Representing nucleotide sequences to neural networks

Whether derived from a double-stranded or single-stranded genome, any nucleotide sequence encodes information in the forward or reverse direction. Conventionally, the sequence is represented by one-hot encoded vectors. For example, adenine (A) could be represented as [1,0,0,0] and thymine (T) could be represented as [0,0,0,1]. One-hot representation of forward and reverse strands results in two L x 4 vectors where L is the length of the sequence. Input vectors to nucleotide-based models have three dimensions (strand, sequence, nucleotide_representation).

We introduce a novel approach to represent genome sequences to neural networks. Precisely, both forward and reverse strands undergo translation across all three reading frames, and in this process, the 20 amino acids and three-stop codons are represented by learnable embedding (LE) vectors with a dimensionality of four, as our experiments showed no improvements on performance upon increasing the embedding length beyond four. The LE vectors are randomly initialized from a uniform distribution [3]. Consequently, a genome sequence of length L is converted into six numerical matrices, each corresponding to a distinct reading frame. Thus, input vectors to amino acid-based models have three dimensions (n_frames, n_codons_in_sequence, dim_LE_vector). As the length of a codon is three nucleotides, the dimensions of the final output are (6, [(L-2)/3], 4). Our results show that this representation technique performs better than conventional one-hot encoded nucleotide vectors (**Table S5**).

#### Neural networks with shared parameters for all reading frames

Since amino acid motifs useful for identifying the origin of genomic sequences can stem from any of the six reading frames, the model must pay equal attention to all of them. Similarly, nucleotide motifs can occur on both forward and reverse strands. Recent work has used parameter-sharing neural network architectures on genomic sequences [4,5] that utilize the same subnetwork on the forward strand and the reverse complement sequences. We adopted this idea for six-frame translated genomic sequences. We implemented parameter sharing by passing all six translated frames of the input genomic sequence through a common sub-network, similar to the implementation in [5]. We summed up feature vectors arising from each reading frame to form a global feature vector (dim=128) that summarizes information from all frames.

#### Initializing neural networks with correct bias

Since the training dataset is imbalanced, setting the bias of the output layer to reflect this imbalance can aid in achieving initial convergence. This can be achieved by the **Eq. S1**

**Eq. S1**

$$f_i = \frac{e^{b_i}}{\sum_j e^{b_j}}$$

Which has the solution

$$b_i = \ln(f_i)$$

This means that a log-transformed class frequency vector can be used to initialize the output layer’s bias correctly.

#### The baseline model (Base)

All the deep learning models we implemented in this study inherit from a modular architecture consisting of three blocks (**Figure 1, Table S5**): representation learner, pattern recognizer, and classifier. The baseline model only consists of the representation learner and the classifier blocks. All the other models are built by adding a pattern recognizer block to the baseline architecture. The representation learner block contains three stacked dilated convolutional layers, with shared parameters as described above. All models can accept a 2^n bp (n=[9..12]) sequence fragment as input and output a score distribution across the four output classes: phage, bacteria, eukaryotic, and archaeal. In our current study, we set n to 11, i.e. an input layer of length 2,048bp.

**Dilated Convolutional Neural Network with Residual Connections (DCNN-Res)**

DCNN-Res (**Figure S4**) is inspired by the ResNet architecture, which is probably the most popular in the field of computer vision [6]. ResNet architecture makes it possible to create deep neural networks using special connections that help information flow more easily through the network. The pattern recognizer consists of a stack of four dilated 1D convolutional blocks with residual connections (DCBRC) which helps the network to learn more complex features. Last, the feature vectors are fed into the logistic regressor comprising two fully connected dense layers.

**DCNN-Res and Attention model (DCNN-Res-Attention)**

The transformer model in this study is inspired by the image transformer [7]. The output of the representation learner is processed by a stack of four residual blocks followed by two transformer encoder layers. The transformer encoder layers adopt a self-attention mechanism to differentially weigh the significance of each part of the input sequence. The regions in the input that highly correlate to the output class will be weighted high compared to the regions that do not correlate to the output.

**DCNN-Res and LSTM model (DCNN-Res-LSTM)**

Long-short-term memory networks are special recurrent neural networks that can learn long-term dependencies in sequence data. Due to their ability to model order dependence, LSTMS have been extensively used for modeling biological sequences [8]; [8,9]. The pattern recognizer here is composed of a stack of four residual blocks, similar to the pattern recognizer of CNN-Res-Attention, followed by a Bi-directional LSTM layer. The residual stacks are deployed to extract complex patterns from the input sequences. The Bi-directional LSTM layer was added to learn the order of patterns extracted by the previous residual block. Here, the residual block outputs a sequence of patterns extracted from the input sequence. The LSTM stack processes each pattern at a time. If a pattern is important, the network will propagate a summary of it to the next time step. This process continues till the end of the sequence. In the end, the LSTM layer returns a feature vector that summarizes all the important patterns and their positions in the input sequence.

**Interpreting Neural Network's Predictions**

To interpret Jaeger's predictions and gain insight into its behavior, we calculated the gradients of the model's output (logits) with respect to the input, which consisted of six-frame translations of nucleotide sequences.

**Eq. S2:**

$$S(frame_n) = \frac{\partial F_c(frame_1, frame_2, frame_3, frame_4, frame_5, frame_6)}{\partial I}$$

Where,

- $F(frame_n)$ is the output of the neural network given the input $frame_n$,
- $c$ represents the class for which the saliency map is calculated (i.e., the predicted class),
- $frame_n$ is the six-frame translated nucleotide sequence, and
- $S(frame_n)$represent the resulting saliency maps.

A custom script was developed to generate and visualize these saliency maps. The maps were normalized, and forward and reverse strand frames were averaged separately into two vectors. To align with the nucleotide sequence, each saliency map was expanded by repeating each value three times.

To investigate the relationship between gradients and gene density on each strand, we randomly selected 50 prokaryotic genomes and 500 phage genomes. Gene density was defined as the total length of all genes on a strand normalized by the strand's length. The correlation between gene density and gradient magnitude was computed using Pearson's r. One-sided t-tests were performed to assess whether the gradient distributions differed significantly when: (1) gene density on the forward strand exceeded that on the reverse strand (hypothesis: mean gradient distribution for the forward strand is greater), and (2) gene density on the reverse strand exceeded that on the forward strand (hypothesis: mean gradient distribution for the reverse strand is greater).

**Table S5.** Validation metrics for different model architectures that were tested in this study.

| Model | Input type | Input length | Precision | Recall |
|---|---|---|---|---|
| **Base** | Amino acid | 1024 bp | 0.70 | 0.82 |
| | | 2048 bp | 0.77 | 0.86 |
| | Nucleotide | 1024 bp | 0.67 | 0.77 |
| | | 2048 bp | 0.70 | 0.80 |
| **DCNN-Res** | Amino acid | 1024 bp | 0.86 | 0.92 |
| | | 2048 bp | 0.91 | 0.96 |
| | Nucleotide | 1024 bp | 0.87 | 0.84 |
| | | 2048 bp | 0.87 | 0.86 |
| **DCNN-Res-LSTM** | Amino acid | 1024 bp | 0.90 | 0.89 |
| | | 2048 bp | 0.92 | 0.94 |
| | Nucleotide | 1024 bp | 0.85 | 0.85 |
| | | 2048 bp | 0.87 | 0.89 |
| **DCNN-Res-Attention** | Amino acid | 1024 bp | 0.78 | 0.87 |
| | | 2048 bp | 0.80 | 0.93 |
| | Nucleotide | 1024 bp | 0.73 | 0.85 |
| | | 2048 bp | 0.77 | 0.89 |

**Table S6.** Module-wise configuration of four neural network architectures tested in this study. Hyperparameters of each layer are denoted within brackets.

| **Block** | **Baseline model** | **DCNN-Res model** | **DCNN-Res-LSTM model** | **DCNN-Res-Attention model** |
|---|---|---|---|---|
| **Representation learner** | 1D convolution *(kernel length=9, dilation=2, kernel=256)*<br>1D MaxPooling *(kernel length=2)*<br>1D convolution *(kernel length=5, dilation=3, kernel=256)*<br>1D MaxPooling *(Kernel length=2)* | | | |
| **Pattern recognizer** | No pattern recognition block | 1D convolution *(kernel length=5, Dilation=2, kernel=256)*<br>1D MaxPooling *(Kernel length=2)*<br>1D convolution *(kernel length=5 Dilation=2 kernel=256)*<br>1D MaxPooling *(Kernel length=2)*<br>**** The above block is stacked five times*** | Bi-directional LSTM *(LSTM cells=128)* | Dot-product Attention |
| | 1D GlobalMaxPooling | | | |
| **classifier** | Linear transform *(Kernel length=128)*<br>Linear transform *(Kernel length=128)*<br>Linear transform *(Kernel length=128)*<br>Softmax activation | | | |

**Table S7.** Jaeger's command line options and their default values

| Option | Description | Default |
|---|---|---|
| **--fsize** | length of the sliding window (value must be $2^n$). | 2048 |
| **--stride** | stride of the sliding window | 2048 |
| **--model** | switch to another deep-learning model | default |
| **--rc** | minium reliability score required to accept predictions. | 0.2 |
| **--batch** | the number of sequence fragments for parallel processing | 96 |
| **--workers** | number of threads | 4 |
| **--getalllogits** | writes window-wise scores to .npy file | 0 |
| **--getsequences** | writes the putative phage-like sequences to fasta file | 0 |

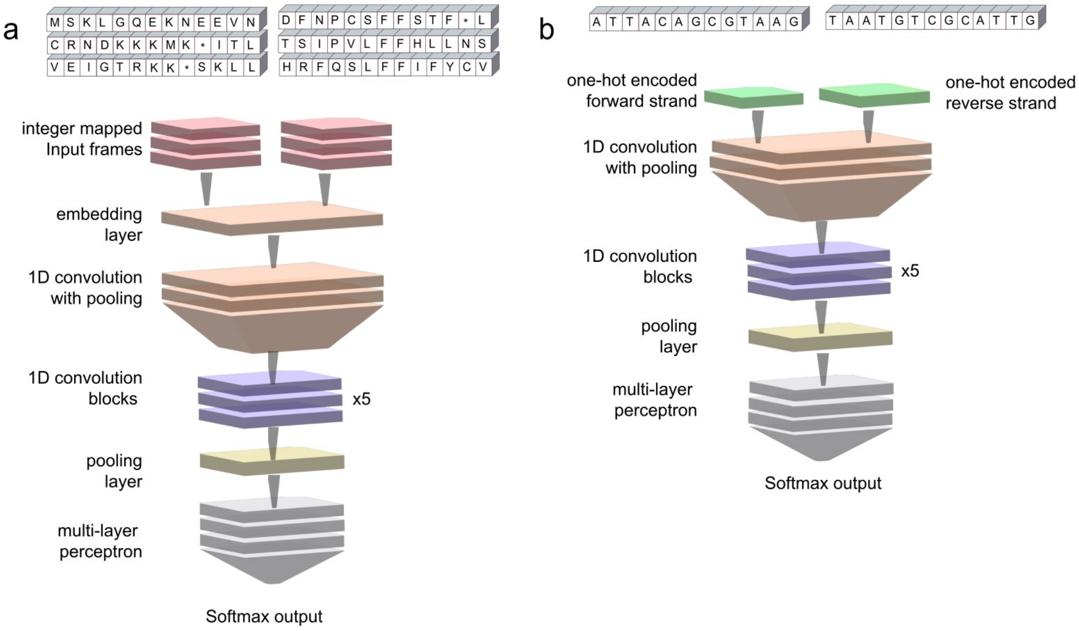


**Figure S4.** Graphical representations of the tested DCNN-Res architecture. **(a)** architecture adapted to operate with the six-frame translated nucleotide sequences. **(b)** architecture adapted to operate with forward and reverse strands. In both cases, the pattern recognizer block was swapped with different layers (*e.g.* Bidirectional LSTM) to obtain other architectures.
