## Supplementary Table 3 for "Jaeger: an accurate and fast deep-learning tool to detect bacteriophage sequences"

**Supplementary table 3.** Results of running Jaeger on 16,702 publicly available assemblies in MGnify.

| | Total | Phage | Non-phage |
|---|---|---|---|
| **Number of contigs >= 2,048bp** | 143,368,698 | 16,152,059 | 127,216,639 |
| **Average length** | 6,912 | 5,300 | 7,116 |
| **The standard deviation of average length** | 14,906 | 8,196 | 15,540 |
| **Min length** | 2,048 | 2,048 | 2,048 |
| **1$^{st}$ quartile length** | 2,533 | 2,473 | 2,542 |
| **Median length** | 3,448 | 3,195 | 3,488 |
| **3$^{rd}$ quartile length** | 5,961 | 4,930 | 6,120 |
| **Max length** | 6,218,175 | 720,900 | 6,218,175 |
| **Contigs >=1Mb** | 875 | 0 | 875 |
