## Supplementary Table 4 for "Jaeger: an accurate and fast deep-learning tool to detect bacteriophage sequences"

**Supplementary Table 4.** Per biome metrics of real-world metagenome benchmark.

| tool | biome | recall | specificity | precision | F1 score | FPR | FDR |
|---|---|---|---|---|---|---|---|
| DeepVirFinder | | 0.29 | 0.92 | 0.48 | 0.34 | 0.08 | 0.52 |
| geNomad | | 0.45 | 0.9 | 0.51 | **0.46** | 0.1 | 0.49 |
| jaeger | | 0.55 | 0.8 | 0.4 | 0.44 | 0.2 | 0.6 |
| PPR-Meta | gut | 0.34 | 0.95 | 0.6 | 0.42 | 0.05 | 0.4 |
| seeker | | 0.19 | 0.84 | 0.27 | 0.2 | 0.16 | 0.73 |
| VIBRANT | | 0.26 | **0.98** | 0.7 | 0.36 | **0.02** | 0.3 |
| VirSorter2 | | 0.29 | **0.98** | **0.75** | 0.41 | **0.02** | **0.25** |
| DeepVirFinder | | 0.71 | 0.92 | 0.75 | 0.73 | 0.08 | 0.25 |
| geNomad | | **0.91** | 0.86 | 0.7 | 0.79 | 0.14 | 0.3 |
| jaeger | | 0.88 | 0.95 | 0.85 | **0.86** | 0.05 | 0.15 |
| PPR-Meta | seawater | 0.9 | 0.78 | 0.59 | 0.71 | 0.22 | 0.41 |
| seeker | | 0.51 | 0.85 | 0.54 | 0.52 | 0.15 | 0.46 |
| VIBRANT | | 0.45 | **0.99** | 0.91 | 0.6 | **0.01** | 0.09 |
| VirSorter2 | | 0.5 | **0.99** | **0.96** | 0.66 | **0.01** | **0.04** |
| DeepVirFinder | | 0.43 | 0.99 | 0.99 | 0.6 | 0.01 | 0.01 |
| geNomad | | **0.88** | 0.98 | 0.99 | **0.93** | 0.02 | 0.01 |
| jaeger | | 0.86 | 0.99 | 0.99 | 0.92 | 0.01 | 0.01 |
| PPR-Meta | soil | 0.86 | 0.92 | 0.96 | 0.91 | 0.08 | 0.04 |
| seeker | | 0.2 | 0.92 | 0.86 | 0.33 | 0.08 | 0.14 |
| VIBRANT | | **0.53** | **1** | **1** | 0.69 | **0** | **0** |
| VirSorter2 | | 0.44 | **1** | **1** | 0.61 | **0** | **0** |
