## Supplementary Figure S1 for "Jaeger: an accurate and fast deep-learning tool to detect bacteriophage sequences"

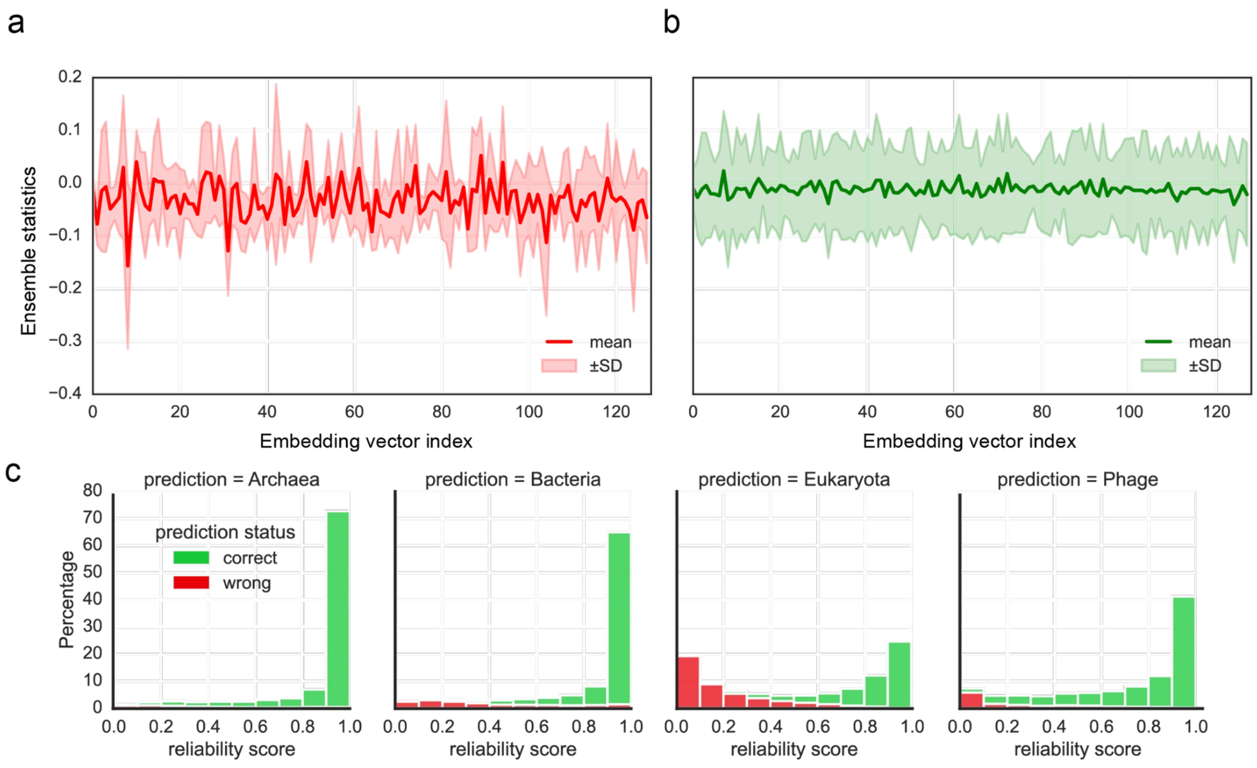


**Supplementary Figure S1.** The reliability module assesses prediction uncertainty by leveraging the penultimate layer embeddings of the neural network. **(a)** The penultimate layer (length=128) embeddings of out-of-distribution (Shuffled genome DB) examples are shown here. **(b)** The penultimate layer embeddings (length=128) of in-distribution (Genome DB) examples are summarised here. **(c)** Summary of the percentage of correct and incorrect predictions per class. Notably, correct predictions had higher reliability scores than incorrect predictions.
