## Supplementary Figure S2 for "Jaeger: an accurate and fast deep-learning tool to detect bacteriophage sequences"

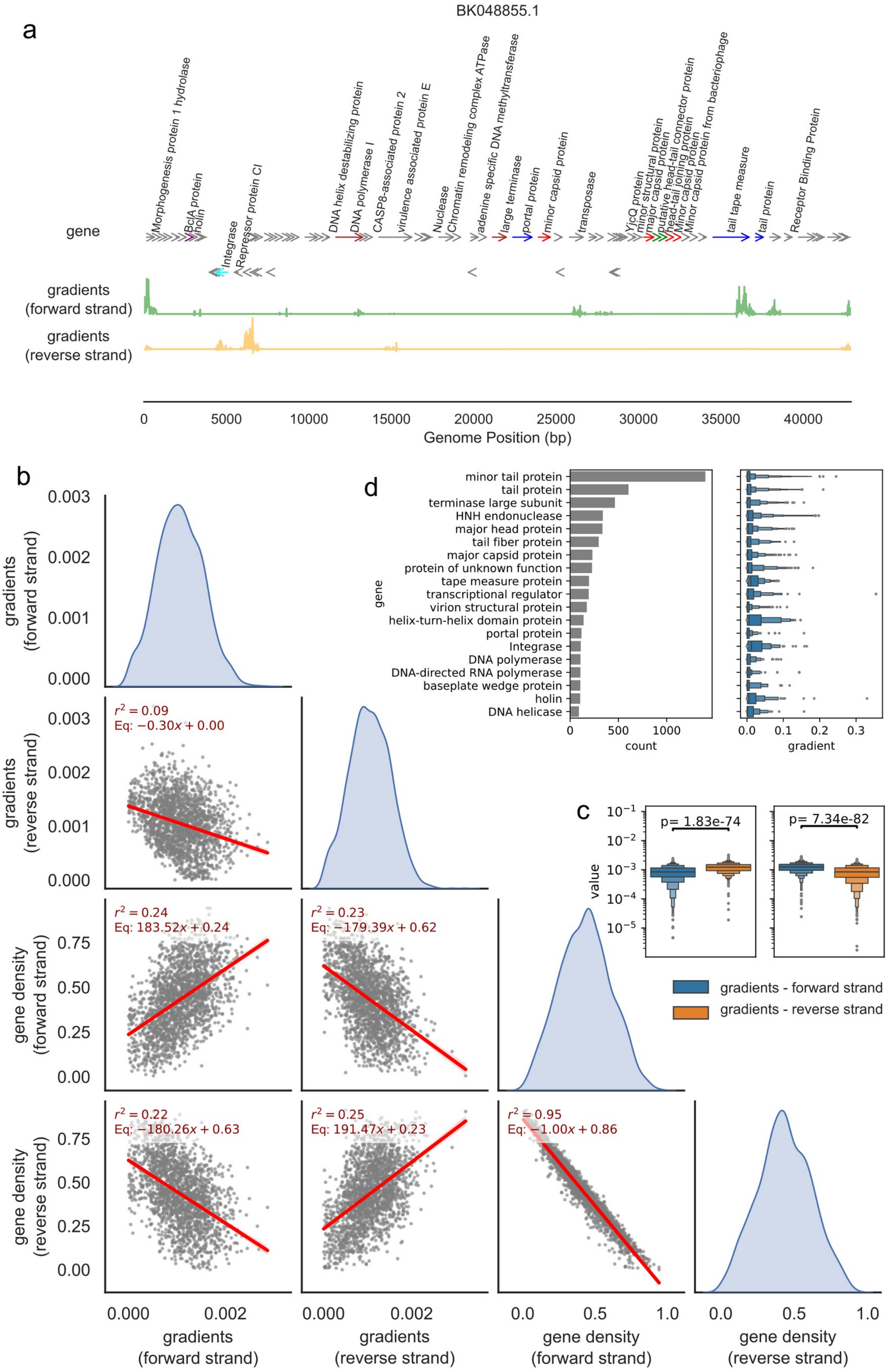


**Supplementary Figure S2.** Analysis of Neural Gradients. (a) The gradients for a partial genome of Caudoviricetes sp. isolate ctU9t19 (BK048855) are displayed. The green line represents gradients for the forward strand, while the green line represents gradients for the reverse strand. Gradients on the forward strand peak at the tail protein, tape measure protein, and morphogenesis protein.

Conversely, gradients on the reverse strand peak at the repressor protein and integrase. This indicates that Jaeger utilizes these specific regions to inform its final predictions. (b) Pair plots illustrate the relationship between gene density on each strand and the magnitude of gradients. There is a positive correlation between gene density and gradient magnitude, suggesting that the model effectively learns to detect genes on either strand. (c) Gradient distributions are shown for two scenarios: (1) when gene density on the forward strand exceeds that on the reverse strand (right) and (2) when gene density on the reverse strand exceeds that on the forward strand (left). Significant statistical differences are observed in the gradient distributions for both cases. (d) The top 20 phage gene categories that exhibit the highest gradient values.
