## Supplementary Figure S3 for "Jaeger: an accurate and fast deep-learning tool to detect bacteriophage sequences"

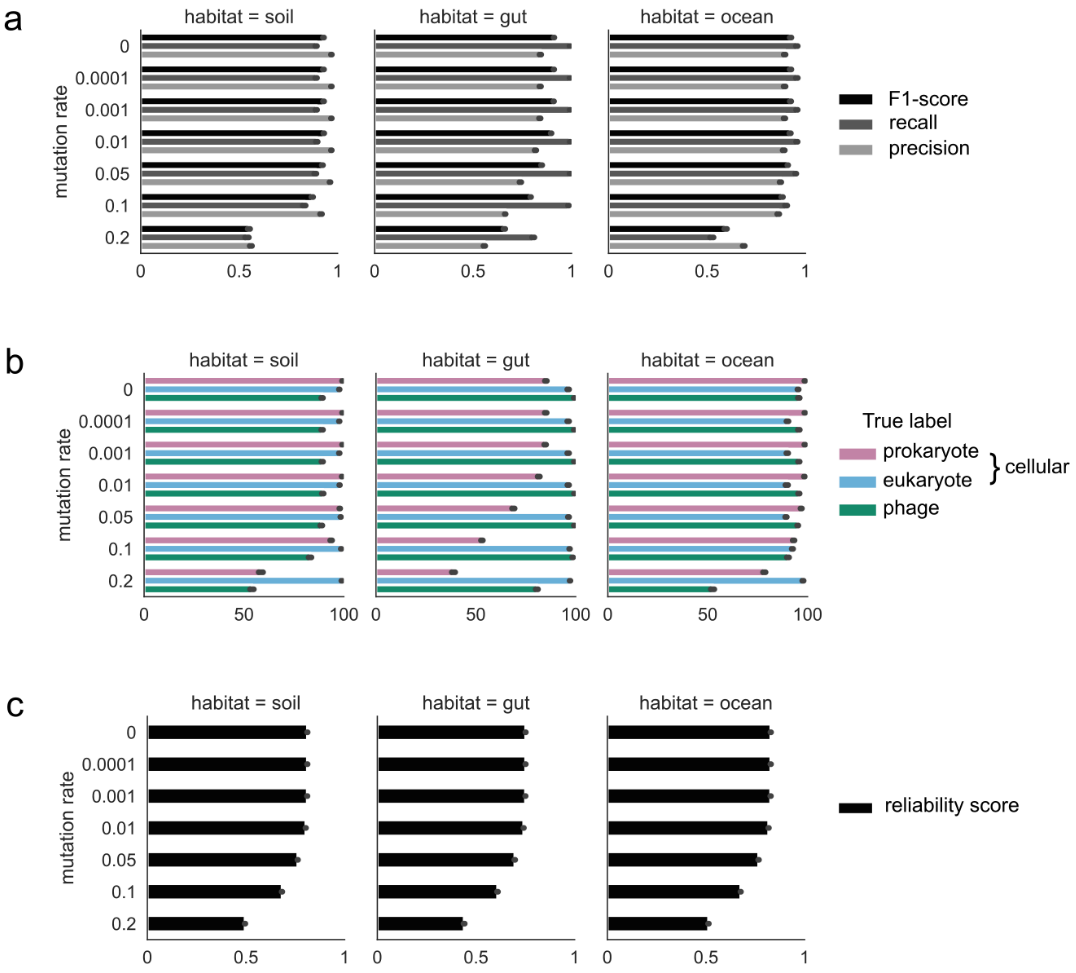


**Supplementary Figure S3.** Genomic sequences from bacteria (n=150), eukaryotes (n=6), and phages (n=150) with biome annotations were selected and fragmented to simulate metagenomic contigs. These fragments were subjected to varying *in silico* mutation rates to assess the impact on predictive accuracy. **(a)** Performance metrics (mean of 10 replicates) across different mutation rates. **(b)** Accuracy at the domain level (mean of 10 replicates) across mutation rates. Green bars represent the proportion of correctly classified phage contigs, while blue and purple bars denote the percentage of eukaryotic and prokaryotic contigs correctly identified as non-phage. **(c)** Influence of mutation rate on the reliability score.
